# T1TAdb: the database of Type I Toxin-Antitoxin systems

**DOI:** 10.1101/2021.04.14.439843

**Authors:** Nicolas J. Tourasse, Fabien Darfeuille

## Abstract

Type I toxin-antitoxin (T1TA) systems constitute a large class of genetic modules with antisense RNA (asRNA)-mediated regulation of gene expression. They are widespread in bacteria and consist of an mRNA coding for a toxic protein and a noncoding asRNA that acts as an antitoxin preventing the synthesis of the toxin by directly basepairing to its cognate mRNA. The co- and post-transcriptional regulation of T1TA systems is intimately linked to RNA sequence and structure, therefore it is essential to have an accurate annotation of the mRNA and asRNA molecules to understand this regulation. However, most T1TA systems have been identified by means of bioinformatic analyses solely based on the toxin protein sequences, and there is no central repository of information on their specific RNA features. Here we present the first database dedicated to type I TA systems, named T1TAdb. It is an open-access web database (https://d-lab.arna.cnrs.fr/t1tadb) with a collection of ~1,900 loci in ~500 bacterial strains in which a toxin-coding sequence has been previously identified. RNA molecules were annotated with a bioinformatic procedure based on key determinants of the mRNA structure and the genetic organization of the T1TA loci. Besides RNA and protein secondary structure predictions, T1TAdb also identifies promoter, ribosome-binding, and mRNA-asRNA interaction sites. It also includes tools for comparative analysis, such as sequence similarity search and computation of structural multiple alignments, which are annotated with covariation information. To our knowledge, T1TAdb represents the largest collection of features, sequences, and structural annotations on this class of genetic modules.

## Introduction

Toxin-antitoxin (TA) systems are encoded within small genetic loci found in most of bacterial genomes including those of pathogens. They are usually composed of two adjacent genes: a stable toxin and a labile antitoxin that inhibits the toxin’s action or expression and whose depletion rapidly leads to cell death or growth arrest (Harms et al. 2018). Six types of TA systems have been described so far depending on the nature and mode of action of the antitoxin (reviewed in (Page and Peti 2016)), and a seventh type has been recently proposed (Wang et al. 2021). While the toxin is always a protein, the antitoxin can be either a protein (types II, IV, V, VI, and VII) or an RNA (types I and III).

Type II TA systems are by far the most well studied class of TA systems. Extensive data for these systems are available in two databases, TADB 2.0 (Xie et al. 2018) and TASmania (Akarsu et al. 2019), which also include limited data for other types of TA systems. These databases also provide a tool (TAfinder and TASer, respectively) to scan and predict TA systems (Akarsu et al. 2019; Xie et al. 2018). Other TA-specific resources include BtoxDB (Barbosa et al. 2015), a database of TA protein structural data, and RASTA_Bacteria (Sevin and Barloy-Hubler 2007), a tool to scan for toxins and antitoxins in bacterial genomes (unfortunately, both are no longer maintained). Overall, none of these existing databases contains expanded data about type I TA (T1TA) systems. The RNA families database Rfam (Kalvari et al. 2021) currently contains more than 600 antitoxin RNA entries that belong to 13 known T1TA families. Rfam is a useful resource that provides, for all families, sequences, alignments, structures, covariance models (that can be used to search genomes), phylogenetic trees, and links to Wikipedia articles. However, it contains no data for the associated toxin mRNAs of the T1TA systems, as Rfam is focused on non-coding RNAs.

A T1TA system consists of a relatively short mRNA (150 to 400 nucleotides long) coding for a small protein (20-60 amino acids in length) whose expression is toxic to the host cell and an antisense RNA (asRNA; 60-200 nt in length) that serves as a counteracting antitoxin to prevent the synthesis of its cognate toxin by directly basepairing to the mRNA. Numerous aspects of T1TA systems have been studied including RNA structure, toxin-antitoxin interaction, regulatory gene expression mechanisms (transcription, translation, degradation, processing), mechanism of action of the toxin, and function in cell physiology (Masachis and Darfeuille, 2018). While TA systems located on plasmids have been demonstrated to contribute to plasmid maintenance (via the mechanism of postsegregational killing, (Greenfield et al. 2000; Gerdes et al. 1986)), the roles of TA systems encoded on the chromosome has only been addressed for a few TA loci (Brielle et al. 2016; Peltier et al. 2020).

A few T1TA systems have been experimentally characterized, but hundreds have been identified by bioinformatic analyses (Arnion et al. 2017; Fozo et al. 2010). By exhaustive amino acid sequence homology searches using PSI-BLAST and TBLASTN run with customized parameters, Fozo and coworkers identified ~900 sequences of ORFs coding for homologs of known type I toxin peptides from 7 families in 95 bacterial species and 229 strains (Suppl. Table S5 in (Fozo et al. 2010)). In addition, through searches based on characteristics of T1TA loci (such as tandem repeats and hydrophobicity) they proposed more than 2,000 novel toxin ORFs in hundreds of genomes (Suppl. Table S8 therein). These analyses were essentially protein-based and did not investigate the mRNA features. Based on thermodynamic local free energies, they could detect the position of asRNA genes of a few known families, but the coordinates spanned only a 100-nt window and the precise start and end boundaries of the full-length molecules were not obtained (Suppl. Table S9 therein). Most of the identified T1TA loci are not yet annotated in genome records, and it is necessary to have an accurate annotation of the complete mRNA and asRNA molecules to understand the co- and post-transcriptional regulation of T1TA systems, which is determined by RNA sequence and structure (Masachis and Darfeuille 2018). Thus, there is a deep need for a central repository of T1TA systems that would include RNA information. In this work, we have built a database, named T1TAdb, which gathers all described and predicted loci of T1TA systems and where the mRNA and asRNA coordinates are annotated. We describe below the content and main features of T1TAdb, along with the procedure used to identify mRNAs and asRNAs.

## Results and Discussion

We have developed T1TAdb, the first database dedicated to T1TA systems. It can be accessed by users through a graphical web interface. A preliminary, development version was put on-line in January 2019 and has been regularly updated, corrected, and improved (https://d-lab.arna.cnrs.fr/t1tadb). The database provides sequence, secondary structure, and genomic information on T1TA loci. In its current, initial version, it is limited to bacterial genomes that were reported in the literature to carry T1TA systems. It contains 1894 loci belonging to 24 families from 218 bacterial species and 493 strains. Data on toxin mRNA, antitoxin asRNA, and toxin peptides were taken from the current literature. Only a small number of loci have been experimentally characterized and the bulk of the data in T1TAdb are mainly based on the results of genome-wide bioinformatic studies. This includes the AapA/IsoA family that has been extensively curated in about 100 genomes of *Helicobacter* and *Campylobacter* (Arnion et al. 2017) and the large number of loci found in hundreds of genomes in the study of (Fozo et al. 2010). The analyses by (Fozo et al. 2010) did not investigate the mRNA genes and recovered only partially the asRNA genes. Therefore, in our work, a major effort to build the T1TAdb database has been devoted to the annotation of the genomic locations of the toxin mRNA and antitoxin asRNA corresponding to each toxin ORF of known family reported by (Fozo et al. 2010), in order to reconstruct the complete loci.

### Annotation of toxin mRNA and antitoxin asRNA genomic localization based on RNA structural features

The genomic regions defining the mRNAs and asRNAs were predicted using RNAMotif (Macke et al. 2001) and RNASurface (Soldatov et al. 2014), respectively. RNAMotif is used to identify regions that can adopt a predefined secondary structure, while RNASurface predicts regions that are structurally more stable than the rest of the genome. A comparison of the asRNA coordinates predicted by RNASurface with those experimentally determined by transcriptome analyses revealed a good agreement. RNASurface predictions were usually within 10 bases of the experimental coordinates, at both the 5’ and 3’ ends, for diverse T1TA families in *Helicobacter pylori* and *Escherichia coli*, but larger differences were more often observed for *Enterococcus faecalis* (Supp. Tables S2-S6). Moreover, the 5’ ends of mRNAs predicted by our RNAMotif structural descriptors showed a similar accuracy. In the case of T1TA systems, a complete TA locus was obtained when a pair of mRNA and asRNA could be predicted, with lengths, orientations, complementary interaction regions, and relative positions matching the genetic organization of a given known TA family (Figure 1 and Suppl. Table S1; see Materials and Methods for details). As can be seen in Table 1, using this procedure, we were able to recover RNA regions of the expected family for 83 to 89% (depending on the family) of the toxin ORFs belonging to five of the seven known families detected by (Fozo et al. 2010) (Suppl. Table S5 therein). The success rate was a bit lower for the TxpA/RatA family (75%), and particularly low for the Fst/RNAII family (29%). For most antitoxin asRNAs of the Fst/RNAII family, RNASurface predicts a sequence that folds into a secondary structure different from the typical RNAII structure and IntaRNA does not identify an interaction region with the toxin mRNA at the expected location. Whether this is a prediction issue specific to this family or if this actually represents structural heterogeneity among Fst/RNAII loci requires further study to be determined. In a few additional cases (less than 10% for each family), a locus could be recovered, but was assigned to a different family. For example, locus TA05747 from *Streptococcus thermophilus* classified in the Fst/RNAII family (based on peptide features) was identified by our procedure (based on RNA and genetic organization features) as a member of the TxpA/RatA family. This discrepancy could be due to a wrong family assignment by either bioinformatic method or peculiarities in the locus that make it fortuitously look more similar to another family. Important discrepancies occurred for 17 loci (from known families) where our procedure assigned a locus from a Gram-positive bacterium to a family found in Gram-negative organisms (or vice-versa) exhibiting a completely different organization. These were clearly erroneous hits and half of them (e.g., TA07190 from *E. coli* and TA07194 from *Salmonella enterica*) were manually corrected and found to match the family predicted by (Fozo et al. 2010). Interestingly, in many of the remaining cases of discrepancy the families show similarities. For instance, locus TA06047 from *E. coli* classified in the Ldr/Rdl family (based on peptide features) was identified as a member of the Hok/Sok family (based on RNA and genetic organization features). However, Ldr/Rdl systems exhibit an organization (including a regulatory leader ORF) similar to that of Hok/Sok systems and may be regulated in the same manner (Kawano 2012), thus the RNA-based classification reflects these similarities. Locus TA05436 from *Staphylococcus aureus* is classified as Fst/RNAII according to peptide features while it was identified as SprA1/SprA1as according to RNA features, but SprA1/SprA1as systems are in fact homologs of Fst/RNAII systems (Weaver et al. 2009; Kwong et al. 2010), and similarly for several loci (e.g., TA05467 from *S. aureus*) classified as either TxpA/RatA or SprG/SprF (Pinel-Marie et al. 2014). The structural descriptors that we used for RNAMotif searches were mainly based on key determinants of the toxin-encoding mRNA secondary structure, in particular Shine-Dalgarno (SD) and complementary anti-SD sequences, as well as terminator stem-loops in the 3’ end, stem-loops in the 5’ end, and regions involved in 5’-3’ long-distance interactions. Because transcription and translation are coupled in bacteria, *cis*-encoded regulatory elements prevent premature translation of the toxin mRNA by sequestering the SD sequence, either co- or post-transcriptionally. Depending on the T1TA system, the anti-SD motif is located either at a short distance upstream of the SD sequence, thus creating a sequestering stem-loop, or near the end of the mRNA occluding the SD sequence via a long-distance base-pairing interaction between the 5’ and 3’ extremities of the mRNA (Masachis and Darfeuille 2018). The overall success of our results demonstrates that taking into account these elements, which are essential for the expression and regulation of T1TA systems, is critical for identifying type I toxin mRNAs.

**Table 1.**
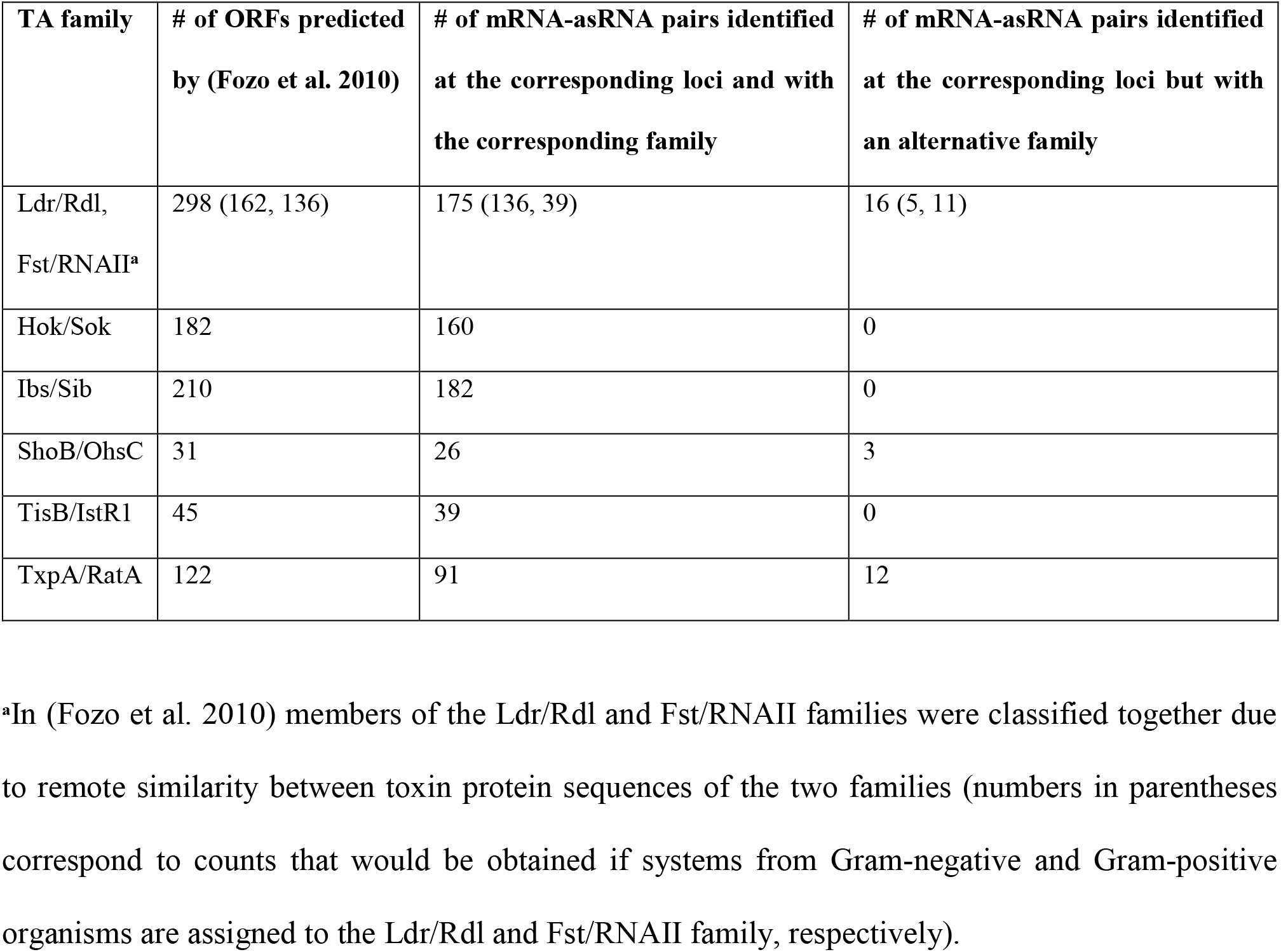
Identification by RNAMotif and RNASurface of toxin-antitoxin mRNA-asRNA pairs corresponding to toxin ORF loci predicted by (Fozo et al. 2010).

| TA family | # of ORFs predicted by (Fozo et al. 2010) | # of mRNA-asRNA pairs identified at the corresponding loci and with the corresponding family | # of mRNA-asRNA pairs identified at the corresponding loci but with an alternative family |
| --- | --- | --- | --- |
| Ldr/Rdl, Fst/RNAlI<sup>a</sup> | 298 (162, 136) | 175 (136, 39) | 16 (5, 11) |
| Hok/Sok | 182 | 160 | 0 |
| Ibs/Sib | 210 | 182 | 0 |
| ShoB/OhsC | 31 | 26 | 3 |
| TisB/IstR1 | 45 | 39 | 0 |
| TxpA/RatA | 122 | 91 | 12 |

<sup>a</sup>In (Fozo et al. 2010) members of the Ldr/Rdl and Fst/RNAlI families were classified together due to remote similarity between toxin protein sequences of the two families (numbers in parentheses correspond to counts that would be obtained if systems from Gram-negative and Gram-positive organisms are assigned to the Ldr/Rdl and Fst/RNAlI family, respectively).

**Fig. 1.**
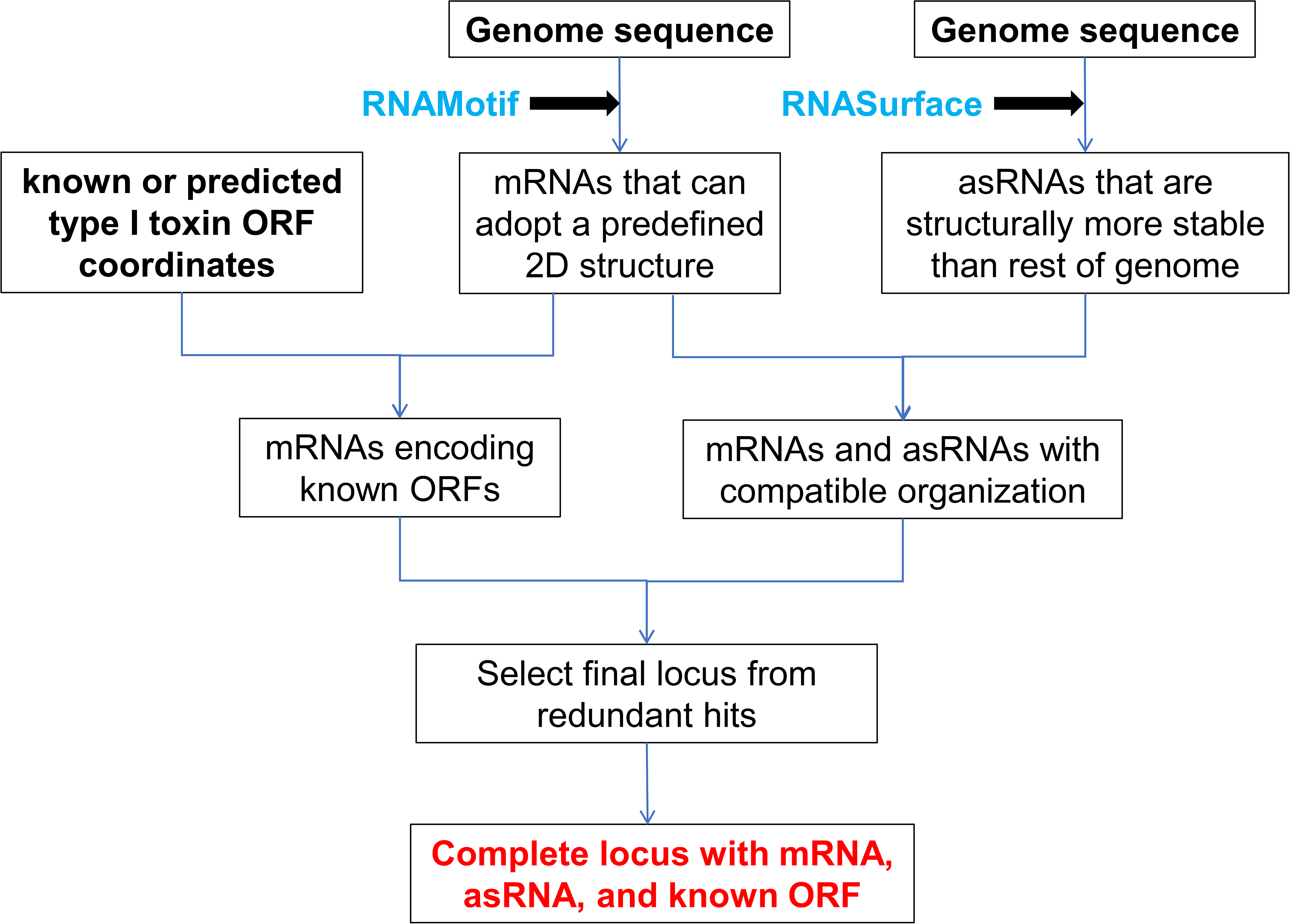
Automatic annotation of mRNAs and asRNAs of known type I TA systems.

### Overview of T1TAdb features and tools

Information in T1TAdb can be searched by entering one or several keywords via the “Keyword Search” tab. A given keyword may be matched against any field in the database or against a specific field such as toxin or antitoxin name, species, strain, taxonomy, locus ID, or genome accession number. Submitting the search form will return a table listing all loci that match the search criteria. The “Select loci” button in the upper left corner of the table allows to further refine the selection of loci by replicon (chromosome or plasmid), TA family or taxonomy, and to select/deselect all loci. The table indicates the locus identifier (TA*nnnnn*), TA family, host strain and taxonomy, and reference publication of each locus. For loci that were predicted using structural descriptors of a TA family different from that reported in the literature (i.e., in (Fozo et al. 2010)), the predicted family is indicated in the “putative family” column, and groups of loci from the same genomic region matching multiple family characteristics appear under a specific identifier (TAG*nnnn*; “Overlapping Locus Group”, see Materials and Methods for details). The leader ORF is also indicated for loci that have been described in the literature to carry a regulatory leader ORF or for which a leader ORF is annotated (for Hok/Sok and Ldr/Rdl families). The loci table provides explanatory notes that appear when hovering over the red asterisk next to specific items. Clicking on a locus identifier leads to a “Locus Details” page that gives detailed information about the locus. The page is divided into four panels (Figure 2). The “Locus” panel provides genomic information with an interactive full genome map (drawn with CGView; (Stothard and Wishart 2005)) of the host strain, showing the location of all TA loci harbored by the strain, and an embedded interactive genome browser (IGV; (Robinson et al. 2011); https://github.com/igvteam/igv.js/wiki) for viewing the genomic context around the locus. The “mRNA” and “sRNA” panels provide the sequence and secondary structure diagram of the toxin mRNA and antitoxin asRNA, respectively. The location of relevant motifs (promoter −10 box, start/stop codon, SD sequence, leader ORF, mRNA-asRNA interaction region) are annotated on the sequences and /or diagrams. The “Peptide” panel shows the sequence, secondary structure model and hydrophobicity plot of the toxin peptide. The “mRNA”, “sRNA”, and “Peptide” panels include a “BLAST Sequence” link for launching a sequence homology search (using BLAST+; (Camacho et al. 2009)) of the selected query against every mRNA, asRNA, ORF, peptide, or genome sequence in T1TAdb. The database also allows the user to input or upload one or several query sequences to perform a BLAST search via the “Sequence Search” tab.

**Fig. 2.**
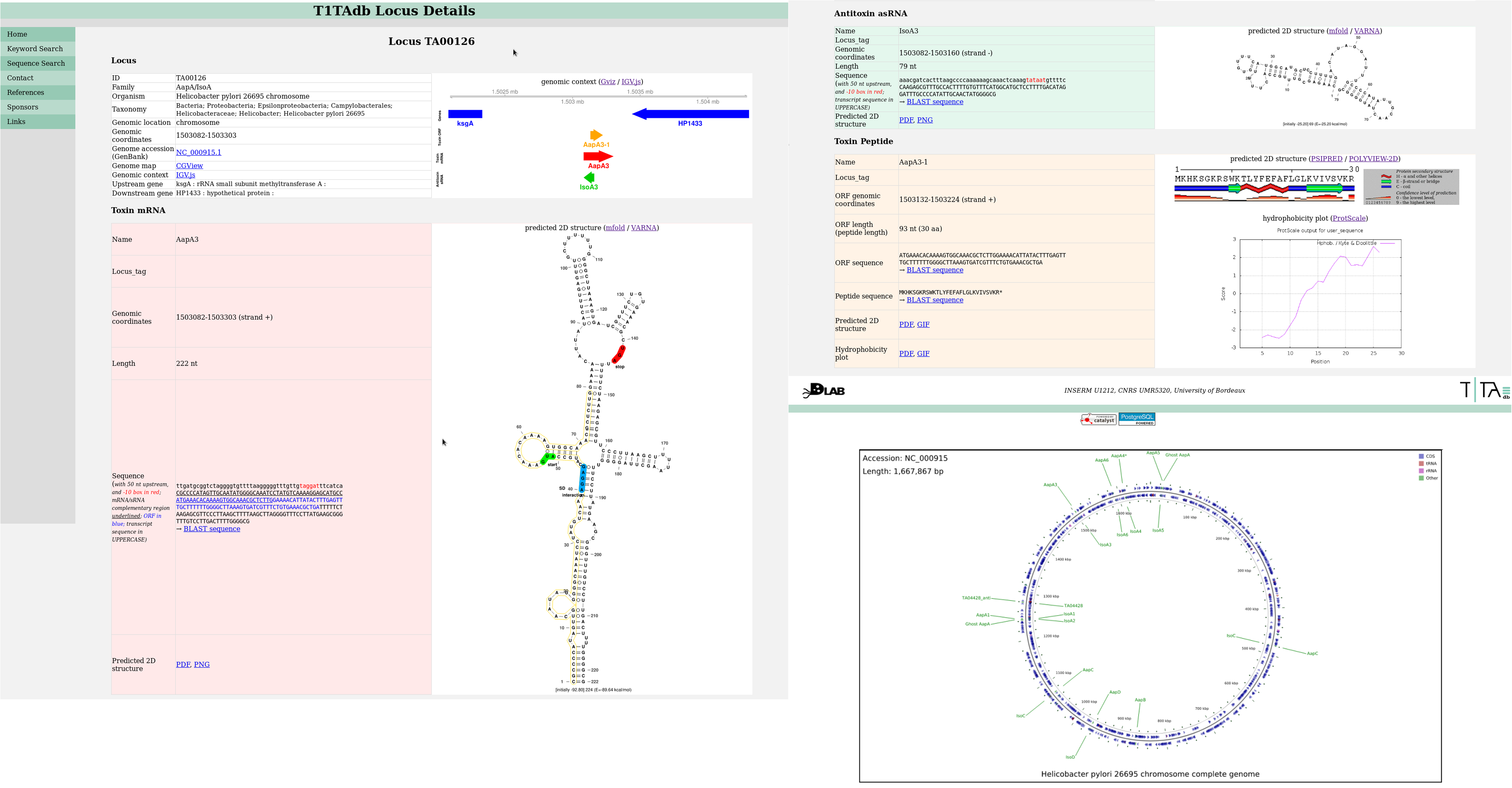
Screenshot of a “Locus Details” page in T1TAdb showing the various panels with sequence, structure, and genomic information.

The T1TAdb home page contains links to a full list of organisms (“view organism list”; “T1TAdb Organisms” page) and TA families (“view family list”; “T1TAdb Families” page) included in the database. Selecting a particular organism name or family name will return a table listing all loci that belong to that organism or family. The family table on the “T1TAdb Families” page provides links to the corresponding Rfam entries for antitoxin RNA families that are included in Rfam. In addition, the table is organized to indicate groups of T1TA families that share some degree of homology (in sequence, structure, and/or genetic organization).

A notable implementation of T1TAdb is that, in addition to homology searches by BLAST, users can perform multiple sequence alignments (using MAFFT;(Katoh and Standley 2013)) to compare mRNA, asRNA, or peptide sequences. Alignments can be launched via the “Align selected sequences” button on top of the table that is returned after selecting loci by organism or family, or following a keyword search. An embedded interactive alignment viewer (MSAViewer; (Yachdav et al. 2016)) is provided to browse and download the alignments. For RNA sequences, structural information is taken into account in MAFFT to produce a structural alignment (Katoh and Toh 2008), and a consensus 2D structure is predicted (using RNAalifold; (Bernhart et al. 2008)) from the alignment. Furthermore, a covariation analysis is performed (using R-chie; (Lai et al. 2012)) to annotate the alignment with covariation information ((Tourasse and Darfeuille 2020); dedicated links are provided to download the consensus and covariation data). With these programs (and BLAST) T1TAdb aims to provide a selected set of tools (run with most default parameters) for comparative sequence and structure analysis, although users may wish to use custom settings or some of the numerous other tools that exist for doing this kind of analysis.

In addition to alignment results, the other data in T1TAdb are also available for download and further custom analysis. Once a set of loci has been selected, the corresponding data can be downloaded using the “Download selected loci” menu at the top of the loci table. Users can obtain a spreadsheet file containing the detailed information about the loci (including host organism, genomic coordinates and sequences of the RNA, ORF, peptide, and promoter features) or sequence files in FASTA format. In the “Locus Details” page for a given locus the structure diagrams can be saved in various formats (SVG, PDF, PNG, or GIF).

The list of publications from which sequences, structures, and other information about the T1TA systems have been obtained is given in the “References” page. Links to other resources on TA systems, as well as the various software and tools used to build T1TAdb, are provided in the “Links” page.

### Example of insights gained from the use of T1TAdb

We present below an example illustrating the usefulness and added value provided by T1TAdb. In particular, we would like to highlight the type of sequence and structure analysis that can be done with the database. Examination of the structural diagrams of individual Sok antitoxin asRNAs from the Hok/Sok family suggested that there may be different structural types. This hypothesis could be tested in T1TAdb by performing structural alignments of different subsets of Sok RNAs which revealed two subgroups: one subgroup had a 2D structure conforming to that reported by (Franch et al. 1997) (Figure 3A), while the other carries an additional stem-loop element at the 5’ end (Figure 3B). The covariation analysis that is run on the alignments in T1TAdb confirmed that this extra element is well supported by covarying positions that reveal compensatory substitutions in a number of sequences. Furthermore, the annotations of the Sok sequences provided in T1TAdb indicated the presence of a −10 box promoter motif (TANNNT, where N means any nucleotide) at an appropriate location upstream of the stem-loop. Although these observations should be verified experimentally, this example shows the potential of T1TAdb to reveal new features in RNA-mediated regulation of T1TA systems by comparative analysis that could not be identified during the study of single T1TA loci.

**Fig. 3.**
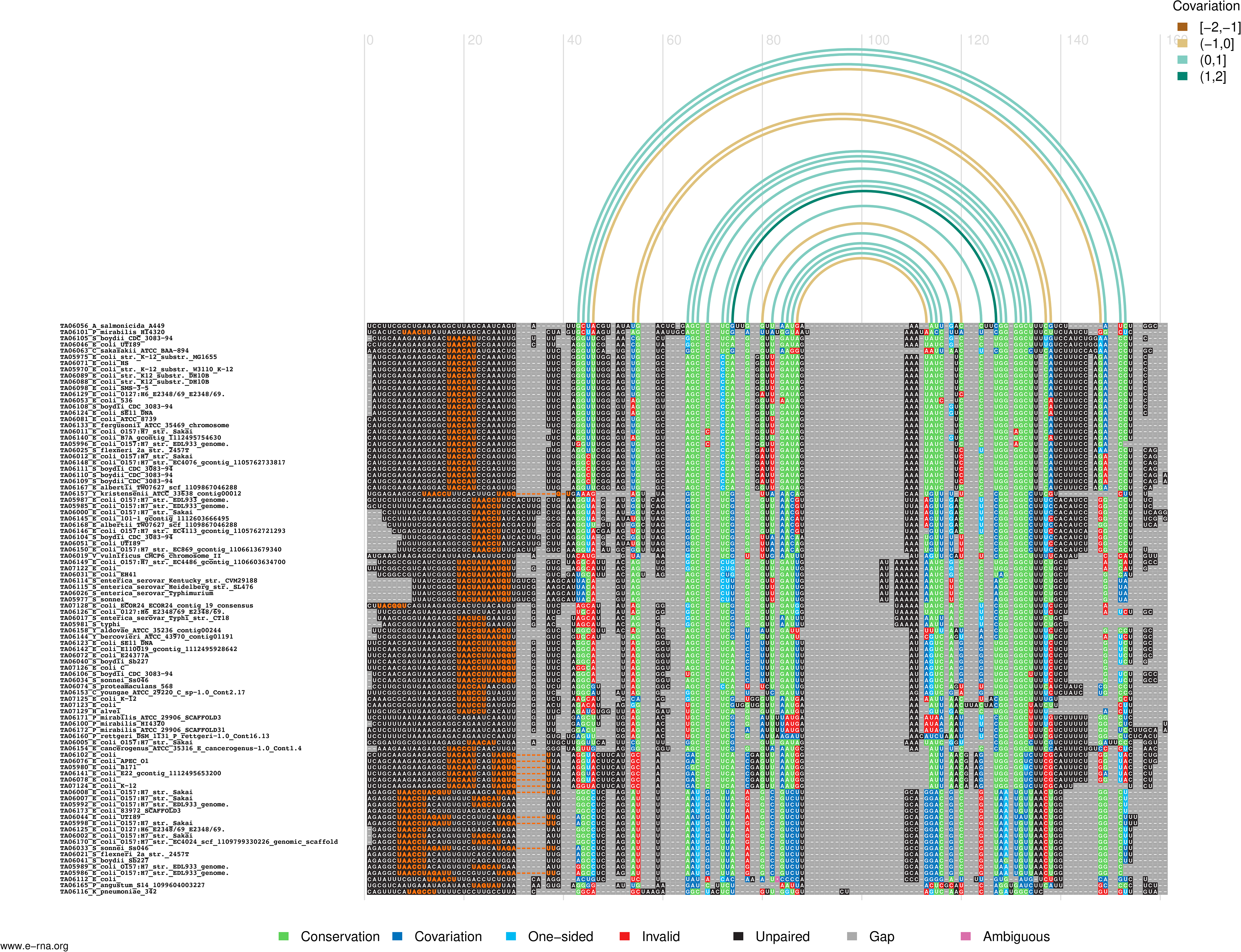
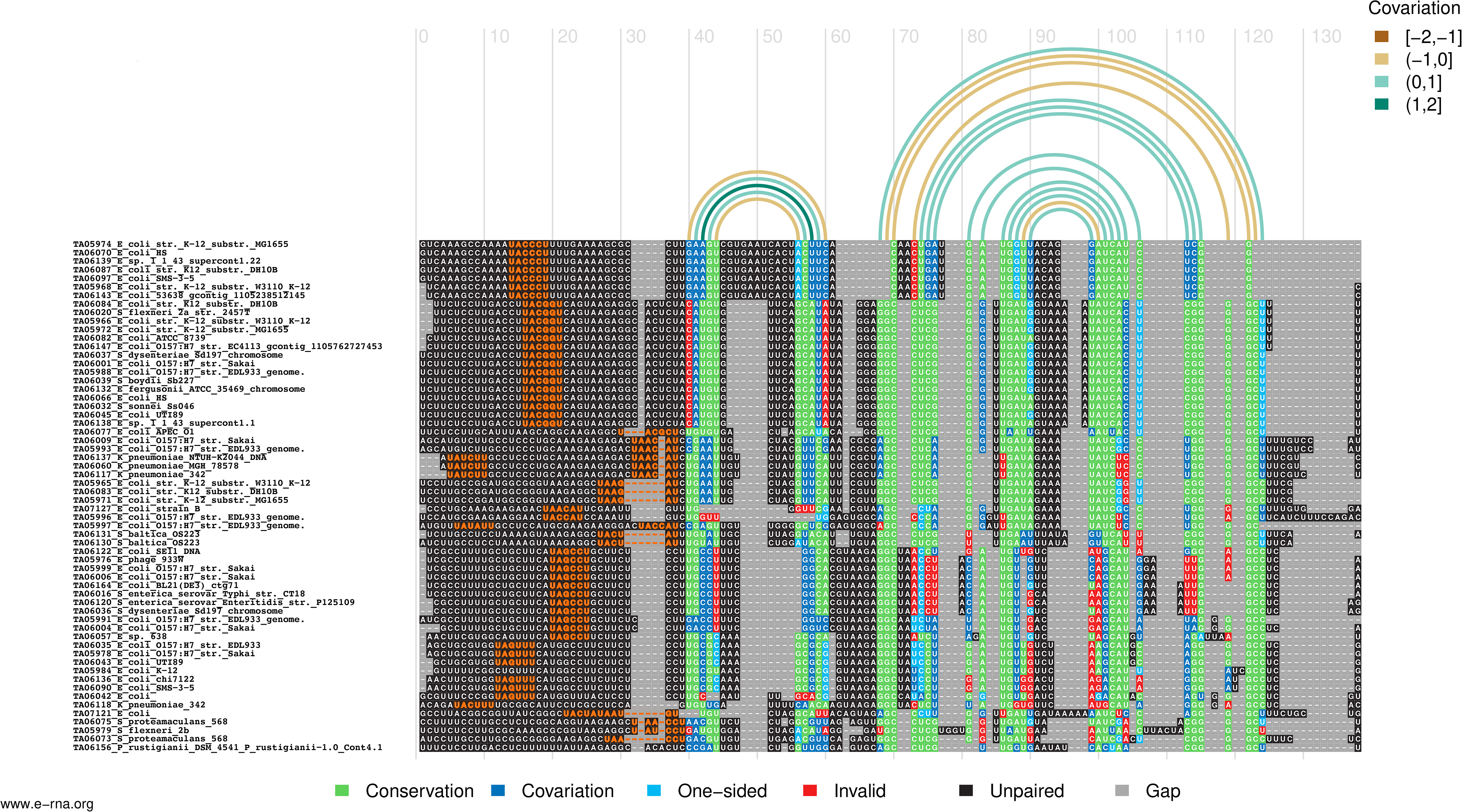
Structural alignments of two subgroups of Sok asRNAs from various Gram-negative bacteria. The structure of the subgroup shown in panel A matches that reported by (Franch et al. 1997) while the subgroup shown in panel B exhibits an additional stem-loop element at the 5’ end. Alignments were computed using MAFFT (and slightly corrected manually) and annotated with covariation information using R-chie. A region of 30 nt upstream of the RNAs was added that includes matches to the −10 promoter box motif (UANNNU, where N means any nucleotide) highlighted in dark orange.

### Conclusions and perspectives

In this work, we have built the first web database totally dedicated to T1TA systems, named T1TAdb. T1TA systems are widespread in bacteria, including pathogenic species, and are studied for numerous aspects including RNA and protein structure and function, regulation of gene expression and cell physiology. In addition to giving access to the collection of loci that have been reported in the literature, the database brings an added value by the annotation of toxin mRNAs and antitoxin asRNAs of loci that were predicted solely by analysis of the toxin peptide sequence, and by providing tools to compare their sequences and structures. This information, together with a wealth of sequence, structure, and genomic data provided on T1TA systems, along with the other databases and tools dedicated to TA systems (such as TADB and TASmania), will certainly serve the scientific community to gain deeper insights of the distribution, evolution, structure, and function of TA systems. This could reveal to be particularly important for the TA systems located in bacterial chromosomes, whose function remains largely unknown. We also believe that the knowledge contained in T1TAdb, and more specifically the RNA structures and loci organization, is a strong asset to help design new tools for identifying new T1TA systems.

In future developments of T1TAdb, we plan to expand the collection of T1TA systems by predicting TA loci in all bacterial genomes available. In addition, the procedure described in this work for the annotation of mRNAs and asRNAs of T1TA systems, which is based on structural features, can be turned into an automatic *de novo* prediction tool, most likely with additional steps to control the false-positive rate. The RNA data in T1TAdb may also be used to generate alignments and covariance models in order to search genome databases for conserved RNAs, as exemplified in the Rfam database (Kalvari et al. 2021). We also anticipate to incorporate transcriptome (RNA-Seq) data into T1TAdb. As the number of sequenced bacterial transcriptomes is increasing, such information would be valuable to check whether TA loci are expressed and to verify the genomic coordinates of the predicted RNAs. Further improvements to T1TAdb will also include tools for simultaneously visualizing and comparing the genomic context (gene neighborhoods) across multiple strains or for the reconstruction of phylogenetic trees, and will allow users to submit data for inclusion in T1TAdb.

### Availability

T1TAdb is freely available on-line at https://d-lab.arna.cnrs.fr/t1tadb.

## Materials and Methods

### Data collection

Sequence, secondary structure, and genomic localization of T1TA loci (including mRNA, asRNA, ORF, and promoter −10 boxes) from the various families were taken from the publications where they were initially discovered and/or characterized (Arnion et al. 2017; Durand et al. 2012; Fozo et al. 2010; Weaver et al. 2009; Maikova et al. 2018; Folli et al. 2017; Kristiansen et al. 2016; Wen and Fozo 2014; Pinel-Marie et al. 2014; Wen et al. 2014; Jahn and Brantl 2013; Weaver 2012; Fozo 2012; Sayed et al. 2012; Han et al. 2010; Sharma et al. 2010; Darfeuille et al. 2007; Pichon and Felden 2005; Kawano et al. 2002; Pedersen and Gerdes 1999; Franch et al. 1997; Masachis and Darfeuille 2018; Meißner et al. 2016; Peltier et al. 2020; Germain-Amiot et al. 2019; Guo et al. 2014; Kawano et al. 2007; Andresen et al. 2020). In most cases, reports were on one or few copies of a given TA system, identified or studied in one or a few bacterial strains. Information on TA systems belonging to the AapA/IsoA family identified by bioinformatic analyses and manual curation in ~100 genomes of *Helicobacter* and *Campylobacter* were taken from (Arnion et al. 2017). (Sharma et al. 2010) reported members of the AapB/IsoB, AapC/IsoC, and AapD/IsoD families in six *Helicobacter* strains. For the sake of consistency and completeness, we located the homologues of these loci in all other *Helicobacter* genomes screened by (Arnion et al. 2017) through sequence homology searches (using BLAST+ 2.2.31; (Camacho et al. 2009)) and multiple sequence alignment. (Kristiansen et al. 2016) identified mRNAs and ORFs belonging to the DinQ/AgrB family in 15 Gram-negative bacteria and we devised a procedure to predict the corresponding asRNAs (see below). The majority of the data in T1TAdb are based on the genome-wide bioinformatic searches by (Fozo et al. 2010) who identified sequences of ORFs coding for type I toxin peptides of known and novel families in hundreds of bacterial strains. For toxins belonging to known TA families, we performed computational analyses to locate the coordinates of the toxin mRNA and antitoxin asRNA in order to identify the complete TA locus corresponding to each reported toxin ORF.

### Prediction of toxin mRNA and antitoxin asRNA genomic localization

Identification of mRNAs and asRNAs was based on the characteristics of known examples. The prediction pipeline is summarized in Fig. 1. Genomic locations of toxin mRNAs were determined using RNAMotif 3.1.1 (Macke et al. 2001). RNAMotif takes as input a descriptor file that contains parameters describing the various elements that make up the secondary structure of a particular RNA (helices, loops, etc. and their respective lengths). Using this descriptor RNAMotif scans genome sequences to find regions that can adopt (i.e., can be folded into) the specified structure. RNAMotif descriptors for mRNAs belonging to 16 known T1TA families were written based on sequences and secondary structures reported in the literature cited above. We did not include every structural element in the descriptors, but rather focused on key determinants such as SD and anti-SD sequences (that sequester the SD motif), location of the ORF within the mRNA, stem-loops in the 5’ end or terminator stem-loops in the 3’ end, and regions implicated in 5’-3’ long-distance interactions (textual summary of RNAMotif parameters given in Suppl. Table S1). The remaining parts of the structure were specified as undefined regions. To avoid descriptors being too specific to a given mRNA instance, lengths and distances between the various elements were usually not set to a defined value but to a min./max. value or a relatively broad range of values. All genomes surveyed by (Fozo et al. 2010) (genome sequences downloaded from NCBI RefSeq; (Haft et al. 2018)) were scanned by RNAMotif with descriptors of all 16 TA families. Hits that spanned the coordinates of the ORFs reported by Fozo *et al*. on the same DNA strand were extracted (by means of the “intersectbed” utility from the BEDTools 2.24.0 package (Quinlan and Hall 2010)). Among those, hits that contained ORFs that were in-frame with the ORFs of Fozo *et al*. (ORFs may not always be identical and could differ slightly in length in some cases depending on the start codon used) and whose ORF and total mRNA length matched the typical range of ORF and mRNA length for a given family were retrieved, whether or not the family matched that reported by Fozo and coworkers.

Antitoxin asRNAs were localized using RNASurface 1.1 (Soldatov et al. 2014), which predicts sequence regions that are structurally more stable than the rest of the genome by local folding of segments up to a predefined size. Genome sequences were scanned with RNASurface (run with options “--winmin 50 --winmax 300 -z −2 -d 500”) to identify structured segments of length 50-300 nt that have a z-score lower than −2 (corresponding to a false-positive rate of 5%, (Soldatov et al. 2014)). For a given TA family, RNASurface segments whose length corresponded to the typical asRNA length for that family (+/− 20%) were retrieved.

Putative promoter −10 boxes were predicted by searching (by means of the PERL module Regexp∷Exhaustive) for the presence of the sequence motif TANNNT (where N means any nucleotide) in a window covering the region −20 to −6 upstream of the identified mRNAs and asRNAs. In case where multiple motifs were present the 3’-most (i.e., the closest to the RNA start) was selected.

In order to find the pair of mRNA and asRNA corresponding to the same TA locus, RNAMotif and RNASurface results were intersected (using “intersectbed” from BEDTools) based on the defined genetic organization of the various T1TA families. In the ShoB/OhsC, TisB/IstR1, Zor/Orz, and DinQ/AgrB families, found in Gram-negative bacteria, the asRNA is located 5’ to the mRNA and does not overlap it, whereas in the Fst/RNAII, TxpA/RatA, YonT/as-YonT, SprA1/SprA1as, SprG/SprF, and BsrE-G-H/as-BsrE-G-H families, all found in Gram-positive bacteria, the asRNA is located on the 3’ side of the mRNA and partially overlaps it (overlaps also the ORF except in the Fst/RNAII family); the asRNA is fully overlapped by the mRNA in the AapA/IsoA, Ibs/Sib, Hok/Sok, and Ldr/Rdl families from Gram-negative organisms, but overlaps the ORF only in the AapA/IsoA and Ibs/Sib families (spans the entire ORF in Ibs/Sib), whereas it is located 5’ of the ORF in the Hok/Sok and Ldr/Rdl families ((Wen and Fozo 2014; Arnion et al. 2017; Pinel-Marie et al. 2014; Durand et al. 2012; Sayed et al. 2012); Suppl. Table S1). For families where the two RNAs do not overlap, a max. distance of 300 nt between the RNAs was allowed (except for the DinQ/AgrB family, see below), and for families where the two RNAs do overlap, min. and max. limits were set for the length of the overlap region based on known examples (Suppl. Table S1). In T1TA systems the mRNA and asRNA share a complementary region of interaction. In families with overlapping RNAs this region generally corresponds to the overlap region, but for the Fst/RNAII and SprA1/SprA1as families interaction occurs outside the overlap sequence (Weaver et al. 2009; Sayed et al. 2012). The software IntaRNA 3.1.0.2 (Mann et al. 2017) was used to find mRNA-asRNA pairs that share a complementary region at the expected location for these two families and to identify regions of interaction for families with non-overlapping RNAs (the interaction region was set to be at least 15 nt long). A specific processing was carried out for the DinQ/AgrB family because in some bacterial strains the locus includes an AgrA ncRNA gene that is homologous to the AgrB asRNA and that can be located in-between DinQ and AgrB. In *Escherichia coli* K-12 substr. MG1655 the sequence of the interaction region in AgrA contains a number of mismatches and it has been shown that AgrA does not bind the DinQ mRNA and that only AgrB, which shares a region almost fully complementary to DinQ, acts as an antitoxin (Kristiansen et al. 2016). Therefore, to accommodate for the possible presence of the AgrA and AgrB paralogs, the max. distance between the mRNA and asRNA was extended from 300 to 500 nt. In cases where two asRNAs were predicted in this region by RNASurface and for both of them a possible sequence of interaction (≥ 15 nt) with the mRNA was identified by IntaRNA, we selected the one for which the interaction was predicted to be the most stable (i.e., had the lowest free energy as computed by IntaRNA). The same procedure was followed to predict the asRNAs corresponding to the DinQ-like loci reported in (Kristiansen et al. 2016) where only mRNAs and ORFs were annotated.

Following the characteristics described above, all pairs of mRNAs and asRNAs that were in the correct orientation and distance and that shared a complementary interaction region were identified for each family. This set of pairs was then intersected with the set of mRNAs that were found to encode ORFs corresponding to those reported by (Fozo et al. 2010), to obtain the pairs that include a known ORF (Fig. 1). For a given locus, there were usually several mRNAs and asRNAs matching these criteria because we used a relatively broad range of values for the parameters describing the length, structure, and organization of the RNAs. Due to the flexibility in the RNAMotif structural descriptors multiple overlapping mRNAs spanning the same ORF were always found, differing by their lengths and a few bases in their start/end coordinates. Among those, the one for which a promoter −10 box could be predicted was selected as the mRNA for the given locus. If a promoter was predicted for multiple mRNAs, or if no promoter was found for any of the mRNAs, then RNA sequences were folded using MFOLD 3.6 (Zuker 1989, 2003) and the minimum free energy (MFE) of the most stable structure of each RNA was normalized by sequence length (adjusted MFE; AMFE) to compare the stability among RNAs. The mRNA that had the smallest AMFE was taken as the mRNA for the locus. The use of AMFE was warranted by the fact that mRNA lengths of the different T1TA families are in the range 200-400 nt where AMFE is almost length-independent (Trotta 2014). For some loci, there were also several possible asRNAs matching the TA family characteristics as RNASurface often predicts multiple RNA segments of different lengths overlapping the same genomic region. Among those, the one for which a promoter −10 box was predicted was selected as the asRNA for the locus. If a promoter was predicted for multiple asRNAs, or if no promoter was found for any of the asRNAs, then the one with the lowest RNASurface z-score was chosen as the putative asRNA for the given locus. Z-score was used here as normalized folding stability measure because asRNA lengths are in the range 60-200 nt where AMFE is length-dependent and thus cannot be reliably used to compare stabilities among RNAs (Trotta 2014).

To verify that the correct asRNA was associated with a given mRNA, an alignment of the antitoxin sequences was made for each family to check the homology and completeness of the sequences. The alignment included only the RNAs whose family corresponded to the family of the ORF in (Fozo et al. 2010), in which case one should expect these sequences to be homologous to each other. Incomplete sequences or sequences whose coordinates were shifted relative to the others were manually corrected. No alignment was done for the mRNAs because they are constrained by the location of the ORFs and thus are normally homologous.

If no suitable mRNA-asRNA pair spanning an ORF identified by (Fozo et al. 2010) could be found, that particular ORF was not included in T1TAdb. However, in cases where no mRNA-asRNA pair corresponding to the same family as that assigned to the ORF in (Fozo et al. 2010) could be identified but the ORF was included in an RNA pair of a different family, the locus was retained and a “putative_family” flag was set to indicate that it matched an alternative family. This flag was also set when ORFs from unknown or novel families were part of loci corresponding to known families. If RNA pairs of multiple TA families spanned the same ORF, only the one matching the family of the ORF was retained. If there were no such pair, then alternative loci were filtered according to the type of organism in which the ORF was encoded, i.e., for a Gram-negative bacterium only the RNA pairs from families found in Gram-negative bacteria were retained, and similarly for Gram-positive organisms. If there were no such pairs, then all alternative mRNA-asRNA pairs covering the ORF were retained. The multiple loci spanning a given ORF were organized into an “Overlapping Locus Group”, which represents a group of loci that have been identified using characteristics of different TA families and that overlap the same genomic region, but we could not determine which one is the real locus in this region.

### Database implementation

T1TAdb is implemented as a relational database in PostgreSQL 10.13 (https://www.postgresql.org/). The database schema was designed following the five rules of normalization (http://www.barrywise.com/2008/01/database-normalization-and-design-techniques/) to avoid data redundancy and inconsistent dependency among tables. The graphical web interface was developed using the PERL Catalyst framework (http://www.catalystframework.org/) to control and manage connections and SQL requests to the database. Graphical design of the web pages was done in dynamic and responsive HTML, JavaScript, and Cascading Style Sheets (CSS) (https://www.w3schools.com/), along with the PERL Template Toolkit 2.26 templating system (http://www.template-toolkit.org/). The T1TAdb website is run via the Apache HTTP 2.4.6 server (https://httpd.apache.org/) under the Linux CentOS 7.8 operating system. Secondary structures of RNA were predicted using MFOLD 3.6 (Zuker 2003, 1989) and annotated diagrams highlighting the location of specific motifs (start/stop codon, SD sequence, interaction region)were generated with VARNA 3.93 (Darty et al. 2009). Secondary structures of toxin peptides were predicted using PSIPRED 4.02 (McGuffin et al. 2000) run with PSI-BLAST 2.2.26 against the UniRef90 protein sequence database (https://www.uniprot.org/help/uniref) and drawn with POLYVIEW-2D (Porollo et al. 2004). Hydrophobicity plots were computed with ProtScale (Gasteiger et al. 2005). Interactive genomic maps in SVG format showing the localizations of TA loci were drawn using CGView (Stothard and Wishart 2005). The embeddable JavaScript/CSS version of the IGV browser ((Robinson et al. 2011); https://github.com/igvteam/igv.js/wiki) was used to provide an interactive visualization of the genomic context flanking TA loci, and SVG images of the genomic context were generated by means of the Gviz package (https://bioconductor.org/packages/release/bioc/html/Gviz.html) in R 3.5.3 ((R Development Core Team 2019); https://www.r-project.org/). Sequence similarity searches in T1TAdb are done with BLAST+ 2.2.31 (Camacho et al. 2009) and multiple sequence alignments are computed by MAFFT 7.407 (Katoh and Standley 2013). For peptide sequences MAFFT is run with the method “mafft-linsi” and the option “--localpair”, whereas for RNA sequences MAFFT is run with the method “mafft-xinsi” and the option “--scarnapair” to incorporate structure information and produce a structural alignment (Katoh and Toh 2008). Alignments are visualized using the embeddable MSAViewer (Yachdav et al. 2016), which is part of the BioJS JavaScript tools (Yachdav et al. 2015). In addition, for RNA alignments, a consensus structure is predicted by means of RNAalifold from the ViennaRNA 2.1.9 package (Lorenz et al. 2011; Bernhart et al. 2008) and a covariation analysis is conducted using R-chie (Lai et al. 2012). Other bioinformatic software such as EMBOSS 6.6.0 (Rice et al. 2000), BioPERL (Stajich et al. 2002), and FASTX-Toolkit 0.0.14 (http://hannonlab.cshl.edu/fastx_toolkit/) were also employed to generate and/or process the data included in T1TAdb.

## Acknowledgements

This work was supported by Institut National de la Santé et de la Recherche Médicale [INSERM U1212], Centre National de la Recherche Scientifique [CNRS UMR5320], Bordeaux University, and Agence Nationale de la Recherche [ANR-12-BSV5-0025-Bactox1, ANR-12-BSV6-0007-asSUPYCO]. The web server hosting T1TAdb is provided by the ODS Web Hosting service of CNRS. We thank Dr. Stéphane Thore and Dr. Sébastien Fribourg for providing additional computing power. Finally, we thank all present and past members of the Darfeuille laboratory for fruitful discussions on this project and Anthony Bugaut, Isabelle Iost, Anaïs Le Rhun, Simon Bonabal, and Olga Soutourina for critical comments on the manuscript.

