## Supplementary Table S1 for "T1TAdb: the database of Type I Toxin-Antitoxin systems"

**Table S1**. Parameters used to automatically annotate various type I TA families.

| TA family | Parameters of RNAmotif for mRNA annotation | mRNA length (nt) | Peptide length (aa) | asRNA length (nt) | Locus organization[e] |
|---|---|---|---|---|---|
| AapA/IsoA[d] | 5' stem-loop (optional): helix >=5 bp, loop 4-20 nt, tail >=10 nt<br>bottom stem (5' part): >=2 or 5 nt<br>5' region: >=5 or 10-50 nt<br>antiSD stem: 8-15 nt containing antiSD motif 2-5 Y[a]<br>SD stem (5' part): undefined length starting with SD motif AGGAG<br>linker SD-start codon[b]: 8-15 nt<br>middle region: 50-200 nt<br>SD stem (3' part): undefined length<br>3' region: 5-50 or 10-50 nt<br>bottom stem (3' part): > =2 or 5 nt | 150-250 | 28-32 | 78 ±20% | asRNA within mRNA, 5' of ORF (with overlap of 10-40 nt) |
| BsrE/as-BsrE | P1 stem (5' part): >=4 nt<br>5' region: 20-40 nt<br>antiSD stem: 3-10 nt containing antiSD motif 2-5 Y[a]<br>SD-antiSD loop: 3-10 nt<br>SD stem: 3-10 nt containing SD motif AGG<br>linker SD-start codon[b]: 8-15 nt<br>3' region: 130-200 nt<br>P1 stem (3' part): >=4 nt<br>P1-SL5 linker: 3-10 nt<br>SL5 stem-loop: helix >=5 bp, loop 3-10 nt | 200-300 | 25-35 | 160 ±20% | asRNA 3' of mRNA and overlaps ORF by 3-30 nt |
| BsrG/as-BsrG | Same as BsrE/as-BsrE, except 3' region of 180-250 nt | 250-350 | 35-45 | 180 ±20% | asRNA 3' of mRNA and overlaps ORF by 3-30 nt |
| BsrH/as-BsrH | Same as BsrE/as-BsrE, except 5' region of 30-50 nt and 3' region of 150-220 nt | 230-300 | 25-35 | 200 ±20% | asRNA 3' of mRNA and overlaps ORF by 3-30 nt |
| DinQ/AgrB | S1 stem (5' part): >=4 nt<br>5' region: 80-130 nt<br>S1 stem (3' part): >=4 nt<br>middle region: 30-80 nt<br>antiSD stem: 3-10 nt containing antiSD motif 2-5 Y[a]<br>SD-antiSD loop: 3-10 nt<br>SD stem: 3-10 nt containing SD motif GGA<br>linker SD-start codon[b]: 8-15 nt<br>3' region: 80-130 nt<br>SL9 stem-loop: helix >=8 bp, loop 3-10 nt<br>3' tail: 6 nt | 300-400 | 25-30 | 85 ±20% | asRNA 5' of mRNA, no overlap |
| Fst/RNAII[d] | 5' region: 8-15 nt<br>antiSD stem: 2-10 or 3-10 nt containing antiSD motif 2-5 Y[a]<br>SD-antiSD loop: 3-10 nt<br>SD stem: 2-10 nt starting with SD motif 2-5 R[c] or 3-10 nt containing SD motif AGG | 170-230 | 30-40 | 65 ±20% | asRNA 3' of mRNA, with an overlap of 3-50 nt (no overlap with ORF) |

| | | | | | |
|---|---|---|---|---|---|
| | linker SD-start codon[b]: 8-20 nt<br>3’ region: 100-120 nt<br>terminator stem-loop: helix >=5 bp, loop 3-10 nt | | | | |
| Hok/Sok[d] | tac_fbi stem (5’ part): 5-10 nt<br>5’ region: 100-200 nt<br>antiSD stem: 2-10 or 5-10 nt starting with antiSD motif 2-5 Y[a]<br>SD-antiSD loop: 5-50 or 40-50 nt<br>SD stem: 2-10 or 5-10 nt starting with SD motif AGG or AGGAG (with 2 mismatches allowed)<br>linker SD-start codon[b]: 5-15 nt<br>3’ region: 180-230 nt<br>tac_fbi stem (3’ part): 5-10 nt | 300-450 | 45-60 | 65 ±20% | asRNA within mRNA, 5’ of ORF (no overlap) |
| Ibs/Sib | 5’ region: 40-60 nt<br>SD stem: 3-10 nt containing SD motif AGG<br>linker SD-start codon[b]: 0-10 nt<br>3’ region: 80-100 nt<br>antiSD stem: 3-10 nt containing antiSD motif 2-5 Y | 130-200 | 17-20 | 140 ±20% | asRNA within mRNA and spans the entire ORF |
| Ldr/Rdl | 5’ stem-loop: helix >=3 bp, loop 3-10 nt<br>5’ region: 120-180 nt (ending with A)<br>antiSD stem: 3-10 nt containing antiSD motif 2-5 Y[a]<br>SD-antiSD loop: 5-15 nt<br>SD stem: 3-10 nt containing SD motif GGNG<br>linker SD-start codon[b]: 5-15 nt<br>3’ region: 130-180 nt<br>terminator stem-loop: helix >=5 bp, loop 3-10 nt | 300-450 | 30-40 | 60 ±20% | asRNA within mRNA, 5’ of ORF (no overlap) |
| ShoB/OhsC | 5’ region: 100-200 nt<br>3’ stem (5’ part): >=6 nt<br>middle region: 15-30 nt<br>antiSD stem: 4-10 nt starting with antiSD motif 2-5 Y[a]<br>SD-antiSD loop: 3-10 nt<br>SD stem: 4-10 nt starting with SD motif AGGA<br>linker SD-start codon[b]: 8-15 nt<br>3’ region: 50-100 nt<br>3’ stem (3’ part): >=6 nt | 200-300 | 20-30 | 65 ±20% | asRNA 5’ of mRNA, no overlap |
| SprA1/SprA1as | 5’ tail: 2 nt<br>antiSD stem: 3-10 nt containing antiSD motif 2-5 Y[a]<br>SD-antiSD loop: 20-40 nt<br>SD stem: 3-10 nt containing SD motif AGG<br>linker SD-start codon[b]: 8-20 nt<br>3’ region: 90-130 nt<br>H6-L6 stem-loop: helix >=8 bp, loop 3-10 nt<br>H6 bottom region: 4 nt | 180-250 | 25-35 | 60 ±20% | asRNA 3’ of mRNA and overlaps ORF by 3-30 nt |
| SprG/SprF[d] | 5’ tail: 70 or 120 nt<br>antiSD stem: 3-10 nt containing antiSD motif 2-5 Y[a]<br>SD-antiSD loop: 3-10 nt | 200-300 | 15-25 or 20-40 | 140 ±20% | asRNA 3’ of mRNA and overlaps ORF by 3-50 nt |

| | | | | | |
|---|---|---|---|---|---|
| | SD stem: 3-10 nt containing SD motif AGG or AGR<br>linker SD-start codon[b]: 8-15 nt<br>3’ region: 50-80 or 60-120 nt<br>3’ tail: 30 or 60 nt | | | | |
| TisB/IstR1 | 5’ stem (5’ part): >=5 nt<br>5’ region: 50-150 nt<br>5’ stem (3’ part): >=5 nt<br>middle region: 50-150 nt<br>antiSD stem: 4-10 nt containing antiSD motif 2-5 Y[a]<br>SD-antiSD loop: 3-10 nt<br>SD stem: 4-10 nt containing SD motif AGGA<br>linker SD-start codon[b]: 8-15 nt<br>3’ region: 50-150 nt<br>3’ stem-loop: helix >=10 bp, loop 3-10 nt | 300-400 | 25-35 | 75 ±20% | asRNA 5’ of mRNA, no overlap |
| TxpA/RatA[d] | 5’ region: 15-25 nt<br>antiSD stem: 3-10 nt containing antiSD motif 2-5 Y[a]<br>SD-antiSD loop: 3-10 nt<br>SD stem: 3-10 nt containing SD motif AGG<br>linker SD-start codon[b]: 8-15 nt<br>3’ region: 150-250 nt<br>terminator stem-loop: helix >=5 bp, loop 3-10 nt | 230-300 | 30-50 or 50-65 | 220 ±20% | asRNA 3’ of mRNA and overlaps ORF by 50-100 nt |
| YonT/as-YonT | promoter region: 10-15 nt (starting with TANNNT)<br>5’ tail: 6-12 nt<br>SD region: 6 nt containing SD motif AGG<br>linker SD-start codon[b]: 8-15 nt<br>3’ region: 170-190 nt<br>3’ tail: 10 nt | 180-250 | 55-65 | 100 ±20% | asRNA 3’ of mRNA and overlaps ORF by 40-80 nt |
| Zor/Orz | 5’ region: 80-120 nt<br>3’ stem (5’ part): >=4 nt<br>middle region: 40-70 nt<br>antiSD stem: 3-10 nt starting with antiSD motif 2-5 Y[a]<br>SD-antiSD loop: 3-10 nt<br>SD stem: 3-10 nt starting with SD motif AGGA<br>linker SD-start codon[b]: 5-15 nt<br>3’ region: 60-100 nt<br>3’ stem (3’ part): >=4 nt<br>3’ tail: 6 nt | 200-300 | 25-35 | 75 ±20% | asRNA 5’ of mRNA, no overlap |

[a]2-5 Y means 2 to 5 C or U (pyrimidine) nucleotides. This was used as antiSD motif for all families.

[b]The start codon was allowed to be ATG, GTG or TTG for all families.

[c]2-5 R means 2 to 5 A or G (purine) nucleotides.

[d]For these families different values were used for some parameters to accommodate different variants.

[e]For families where the mRNA and asRNA overlap the limits imposed on the length of the overlap region are indicated, and for families where the two RNAs do not overlap the max. distance between the RNAs was set to 300 nt, except for the DinQ/AgrB family where it was set to 500 nt (to accommodate for the presence of AgrA and AgrB paralogs).
